# Markov Katana: a Novel Method for Bayesian Resampling of Parameter Space Applied to Phylogenetic Trees

**DOI:** 10.1101/250951

**Authors:** Stephen T. Pollard, Kenji Fukushima, Zhengyuan O. Wang, Todd A. Castoe, David D. Pollock

**Author notes:** Correspondence to, 12801 E 17^th^ Ave, MS 8101, Aurora, CO 80045.

## Abstract

Phylogenetic inference requires a means to search phylogenetic tree space. This is usually achieved using progressive algorithms that propose and test small alterations in the current tree topology and branch lengths. Current programs search tree topology space using branch-swapping algorithms, but proposals do not discriminate well between swaps likely to succeed or fail. When applied to datasets with many taxa, the huge number of possible topologies slows these programs dramatically. To overcome this, we developed a statistical approach for proposal generation in Bayesian analysis, and evaluated its applicability for the problem of searching phylogenetic tree space. The general idea of the approach, which we call ‘Markov katana’, is to make proposals based on a heuristic algorithm using bootstrapped subsets of the data. Such proposals induce an unintended sampling distribution that must be determined and removed to generate posterior estimates, but the cost of this extra step can in principle be small compared to the added value of more efficient parameter exploration in Markov chain Monte Carlo analyses. Our prototype application uses the simple neighbor-joining distance heuristic on data subsets to propose new reasonably likely phylogenetic trees (including topologies and branch lengths). The evolutionary model used to generate distances in our prototype was far simpler than the more complex model used to evaluate the likelihood of phylogenies based on the full dataset. This prototype implementation indicates that the Markov katana approach could be easily incorporated into existing phylogenetic search programs and may prove a useful alternative in conjunction with existing methods. The general features of this statistical approach may also prove useful in disciplines other than phylogenetics. We demonstrate that this method can be used to efficiently estimate a Bayesian posterior.

## INTRODUCTION

Phylogenetic inference has long played a pivotal role in molecular evolution and evolutionary genomics (e.g. Felsenstein 2004; Vonk 2013; Fukushima 2017). It provides unique information about gene and protein interactions (Wang 2005; Hackett 2007; Reyes-Prieto 2007; Craig 2007) and is critical for detecting adaptive bursts and functional divergence (e.g. Castoe 2008; Castoe 2009). Despite its importance, phylogenetic inference is difficult partly because searching tree space is an NP-hard problem (Bodlaender 1992; Brocchieri 2001). Distance-based methods such as neighbor-joining (NJ; (Saitou 1987)) are fast and often provide good approximate results but are considered less reliable than the computationally expensive (Hershkovitz 1998; Takahashi 2000; Whelan 2001) likelihood-based methods (maximum likelihood, ML, and Bayesian or posterior probability, PP). While distance methods generate a single tree using heuristic approaches, likelihood methods must search tree space, generally by running an optimization scheme or Markov chain Monte Carlo (MCMC). Tree space is often searched using various forms of branch swapping (Felsenstein 1981; Huelsenbeck 1997, 2001; Sullivan 2005; Anisimova 2006). A cautious approach to interpreting results from traditional branch-swapping algorithms is warranted, particularly for trees with sequences from many taxa (Mossel 2005).

The principle confounding effect in phylogenetic inference is that multiple substitutions may occur at the same site. Distance-based methods are inferior to likelihood-based methods in accurately inferring multiple substitutions (Felsenstein 1984; Huelsenbeck 1996; Xia 2006). Distance-based methods are also far more strongly biased by long-branch attraction and cannot fully incorporate the advantages of site-specific models of evolution (Huelsenbeck 1995, 1997; Pollock 1998). Another major class of phylogenetic analysis, based on the principle of maximum parsimony, will not be considered here because parsimony methods are far slower than distance methods, and they do not accurately model evolutionary processes despite having the same biases and inaccuracies as distance methods. The computational limitations of likelihood-based methods become far more severe with large amounts of sequence data from highly diverse sets of organisms (Pollock 2000; Sanderson 2003; A. J. de Koning 2010). For example there are 2.75*10^76^ possible topologies relating 50 taxa (Felsenstein 2004), making exhaustive approaches impossible. Branch-and-bound searches can reduce the tree space to be examined for smaller trees but are insufficient for large datasets because the number of tree topologies is still too large (Hendy 1982). Thus, heuristic searches must be used for large trees, evaluating trees that are proximal to reasonably likely trees that have already been found. These searches are currently often performed using branch-swapping algorithms such as nearest-neighbor interchange (NNI), subtree pruning and regrafting (SPR) and tree bisection and reconnection (TBR) (e.g. Ronquist 2003; Salemi 2009). The number of NNI, SPR, and TBR neighbors of any topology increase respectively as linear, quadratic, and cubic functions of the number of taxa, and the trees proposed are not necessarily of similar likelihood to the known tree. Therefore, many highly improbable trees are evaluated in branch-swapping algorithms, and the correct solution is not guaranteed due to the presence of local optima in tree space (Mossel 2005). Branch length optimization (or posterior equilibration) must also be performed after branch swapping and is an additional source of computational cost.

Several heuristic approaches have been developed to release tree searches from local optima. Ratchet methods employ multiple initial trees perturbed by bootstrap resampling to ensure a less-overlapping tree space in subsequent optimizations using branch swapping (Nixon 1999; Vos 2003). The partial stepwise addition (PSA) approach enables escape from local optima by removing some taxa during the topology search (Whelan 2007). Simulated annealing(SA; Kirkpatrick 1983) and Metropolis-coupled Markov chain Monte Carlo (MCMCMC; Geyer 1991) manipulate a likely range of proposed tree acceptances in a single heuristic search or in multiple interacting chains, respectively. Genetic algorithms (GAs) simulate the population dynamics of tree topologies using likelihood as a fitness parameter (Matsuda 1995). These methods outperform simple heuristic searches in at least some contexts. All approaches listed above employ branch swapping to explore tree space and therefore suffer from inefficiency due to the decoupling of topology proposals from the likelihoods of the topologies.

Here we consider whether the beneficial features of Bayesian analyses under relatively complex models can be profitably combined with the speed of distance methods based on relatively simple models. The key to our approach is that rather than using branch swapping to explore phylogenetic tree space, distance-based trees predicted from partially sampled sequences are used. We use Markov chain Monte Carlo (MCMC) and a Metropolis-Hastings algorithm in which new steps in the chain are proposed based on bootstrap resampling a proportion of the current sequence sample. Heuristic phylogenetic trees based on the new sample are created using NJ and the likelihoods of the new trees are evaluated using the full sequence dataset and the mtMam model (Yang 1998). The unwanted sampling distribution induced by the NJ proposal mechanism is estimated by running the proposal mechanism without calculating the likelihoods of the proposed trees. The posterior is then corrected for this sampling distribution. We evaluated the effect of different site sample sizes used to generate the NJ trees (sample size) and different resample proportions (jump size).

## MATERIALS AND METHODS

### Mitochondrial Sequences

The 495 amino acid COI 1249-taxon mitochondrial gene alignment from Goldstein et al was used (Goldstein 2015). 10 taxa were arbitrarily selected from the alignment to use for testing and are shown in Table 1

**Table 1.**
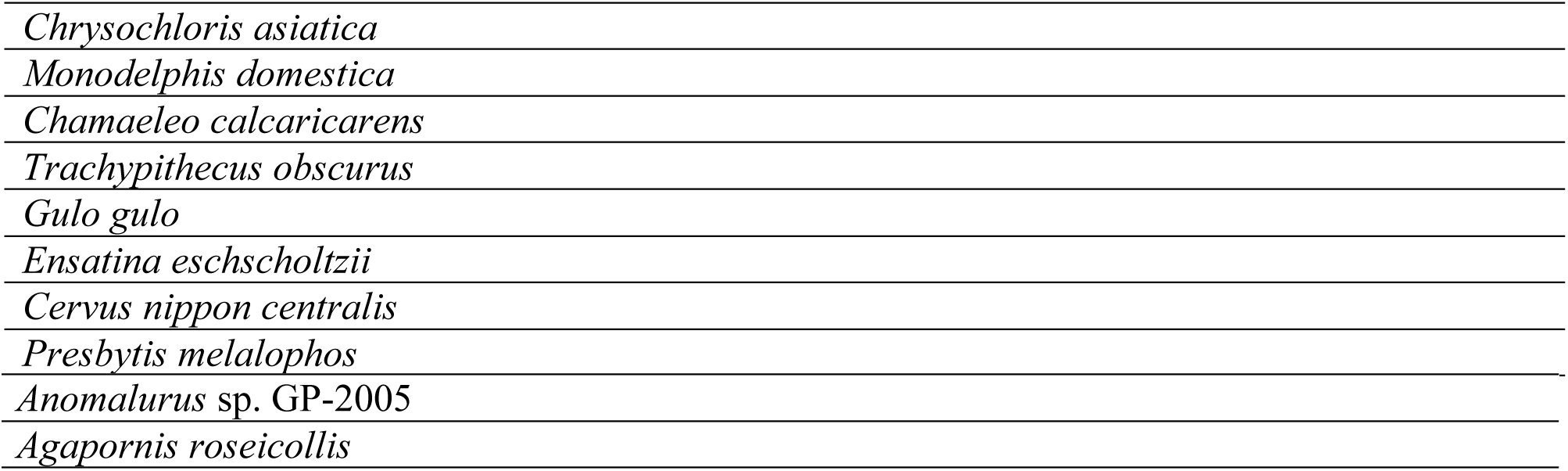
Species in the Cytochrome C Oxidase Subunit 1 Alignment

### Program Details

A Perl program, *MarkovKatana,* was written to implement the Markov chain bootstrapping algorithm. *MarkovKatana* takes multiple sequence alignments in the fasta format and outputs phylogenetic trees in the Newick format, along with likelihood values. Another program *Forest* was written to analyze the trees generated by *MarkovKatana* to calculate tree and branch frequencies. *MarkovKatana* and *Forest* were tested on and are compatible with current Unix-based operating systems as well as Windows. The program *PAML* was used to calculate the likelihoods for the trees using the entire alignment of 495 amino acids (Yang 2007).

### Branch Prior Calculations

Branch priors were calculated as

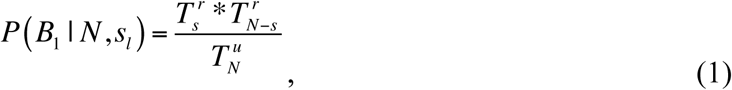

where 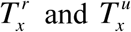 are respectively the number of possible rooted and unrooted topologies with *x* taxa, *N* is the total number of taxa being evaluated, and *s* is the smaller number of taxa that are segregated on one side or the other of branch *B*_*b*_ (Pickett 2005).

### Modifying Implementation of NJ in Markov Katana to Improve Branch Length Estimation

In initial runs, the NJ algorithm often generated unrealistically short branches, so to counteract this we lengthened the shortest branches by adding a random number from 0 to 2 substitutions (a branch length increase of 0 to 2/495). This limited the effect of these implausibly short branches in the proposal mechanism. Short branches were still possible, but extremely short branches were not as likely to be proposed.

## RESULTS

### Details of the Markov Katana Implementation

A bootstrap sampling procedure (Felsenstein 1985; Zharkikh 1995) was employed to sample sites in the alignment that were then used to calculate distance matrices. Although complete and partial bootstrapping has been used extensively in phylogenetic studies to evaluate branch support and tree confidence (Efron 1996; Alfaro 2003), we used it solely to generate a broad distribution of reasonably likely trees based on the NJ heuristic. Note that while partial sampling is more common when employing the related jackknife approach, bootstrapping approaches such as that employed here sample with replacement, rather than without replacement as in the jackknife. Depending on the number of sites sampled (the sample size), trees produced from partial sequence samples can be quite different from the ML tree of the entire alignment and considerably less likely (Castoe 2009). Evaluating the posterior distribution with an importance sampling approach using these trees is not feasible because the extreme variation in likelihoods among trees means that a few trees would dominate the weightedimportance sampling average (Kuhner 1995). Instead, it is necessary to use a progressive Markov chain approach to evaluate the posterior, such as the Metropolis-Hastings algorithm (Hastings 1970), in which the proposed sample depends on the current sample (Fig. 1). Only a fraction of sites is resampled in each generation of the chain. The NJ tree generated from the proposed sample updates both branch lengths and topology simultaneously, and the likelihood of this proposal was then calculated on the full alignment. The number of sites resampled was uniform randomly chosen up to some maximum, which we will call the ‘jump size’.

**Figure 1.**
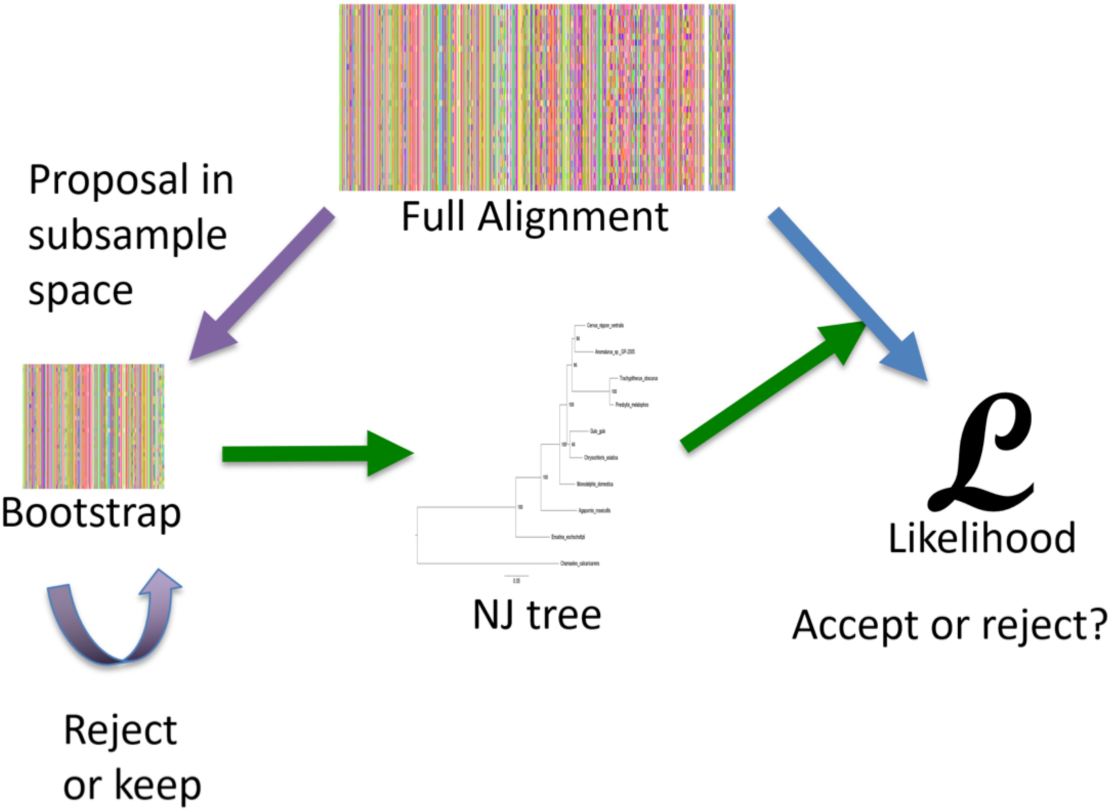
Flow of the Markov katana procedure.

### Posterior Calculations

To obtain the posterior, the uncorrected distribution of trees after the initial Markov katana (MK) run must be corrected for the bias induced by the proposal mechanism. In these runs, the sample size as a fraction, *f*, and the jump size, *j*, were variable parameters and differed among runs as specified. For a given sampled generation, *k*, the alignment sample at that generation produced a NJ genealogy, *G*_*k*_, with topology, *T*_*i*_. The proportion of times that each different topology was produced by the chain out of *K* sampled generations in the chain is an estimator of the uncorrected posterior for a given sample size, *f*, or

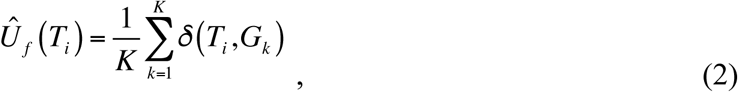

where 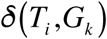 is a delta function equal to 1 if _*G*__*k*_ has topology *T*_*i*_ and otherwise 0. To obtain the corrected topology posterior, *C*(*T*_*i*_), we first estimate the topology sampling bias 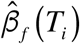induced by NJ proposals with sampling fraction *f*, by sampling *K’* genealogies from a separate chain in which all proposals are accepted, to obtain

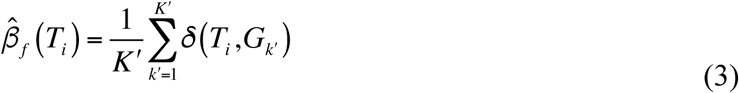

We note that this procedure is identical to obtaining a NJ partial bootstrap, but by running the Markov chain with a given jump size we can obtain the connectedness among topologies, providing a natural topological distance measure.

We then recognize that

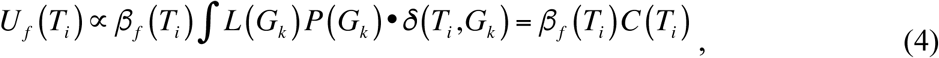

where *L*(*G*_*k*_), *p*_*f*_ (*G*_*k*_), and *P*(*G*_*k*_), are the likelihood, the genealogy sampling bias induced by NJ proposals with sampling fraction *f*, and the prior, respectively. Here we assume a flat prior across all tree topologies. The next step is to divide the uncorrected topology posterior by the sampling distribution induced by the proposals to obtain

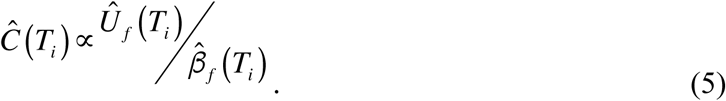

We normalized the corrected posteriors by dividing by the sum of all corrected posteriors over all topologies sampled.

It may sometimes be useful and possibly more accurate to calculate branch (a.k.a. aspecies bi-partition, or edge) posteriors directly over the sample of trees,

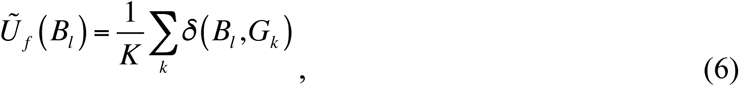

where *δ*(*B*_*l*_,*G*_*k*_) is a delta function equal to 1 if _*G*__*k*_ has branch *B*_*l*_, and otherwise 0. The ~ symbol indicates that the branch uncorrected posteriors were calculated directly. In this case, it is necessary to appropriately adjust for the sampling distribution on the branch induced by ! and a similarly topological constraints (Pickett 2005), which is contained in both *Ũ*_*f*_ (*B*_*l*_) and a similarly obtained

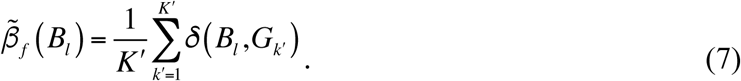

This prior is put back into the posterior calculation as

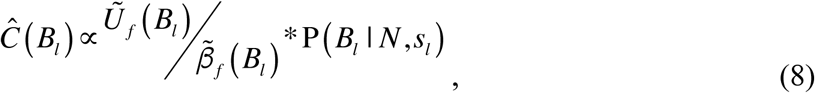

where P (*B*_*l*_ | *N*,*s*_*l*_) is the prior probability of branch *B*_*l*_ induced by topological structures, N is the total number of extant species in the tree, and *s*_*l*_ is the smaller number of species that are partitioned to one side of branch *B*_*l*_. P(*B*_*l*_ | *N*, *s*_*l*_) can be calculated directly (see Methods).

### Implementation of the Markov katana

We began by analyzing a 10-taxon Cytochrome C Oxidase subunit 1 (COI) amino acid alignment (495 residues) that was chosen so that there would be a moderate level of topological uncertainty in the posterior (Fig. 2). Preliminary evaluations indicated that NJ trees on bootstrapped data have a distribution of topologies that are relatively similar among distance types (Sup. Fig. S1). Although there is considerable noise to the estimates for very small frequencies, and there is a slight shift towards higher frequencies with the Markov katana difference NJ, overall the two measures have a nearly linear relationship. This gave us confidence that NJ trees based on differences rather than corrected distances might be sufficiently accurate for our purposes, so to keep the NJ calculations as simple and fast as possible for initial testing, distances were generated using the simple difference matrix. The likelihood of the proposed tree topology and branch lengths were then evaluated using the mtMam substitution rate model (Yang 1998) on the entire sequences. Continuing to keep things simple for initial testing, we used a flat prior, although we imagine that most future implementations will want to incorporate other priors here, such as the commonly used exponential priors on branch lengths (Yang 2005).

**Figure 2.**
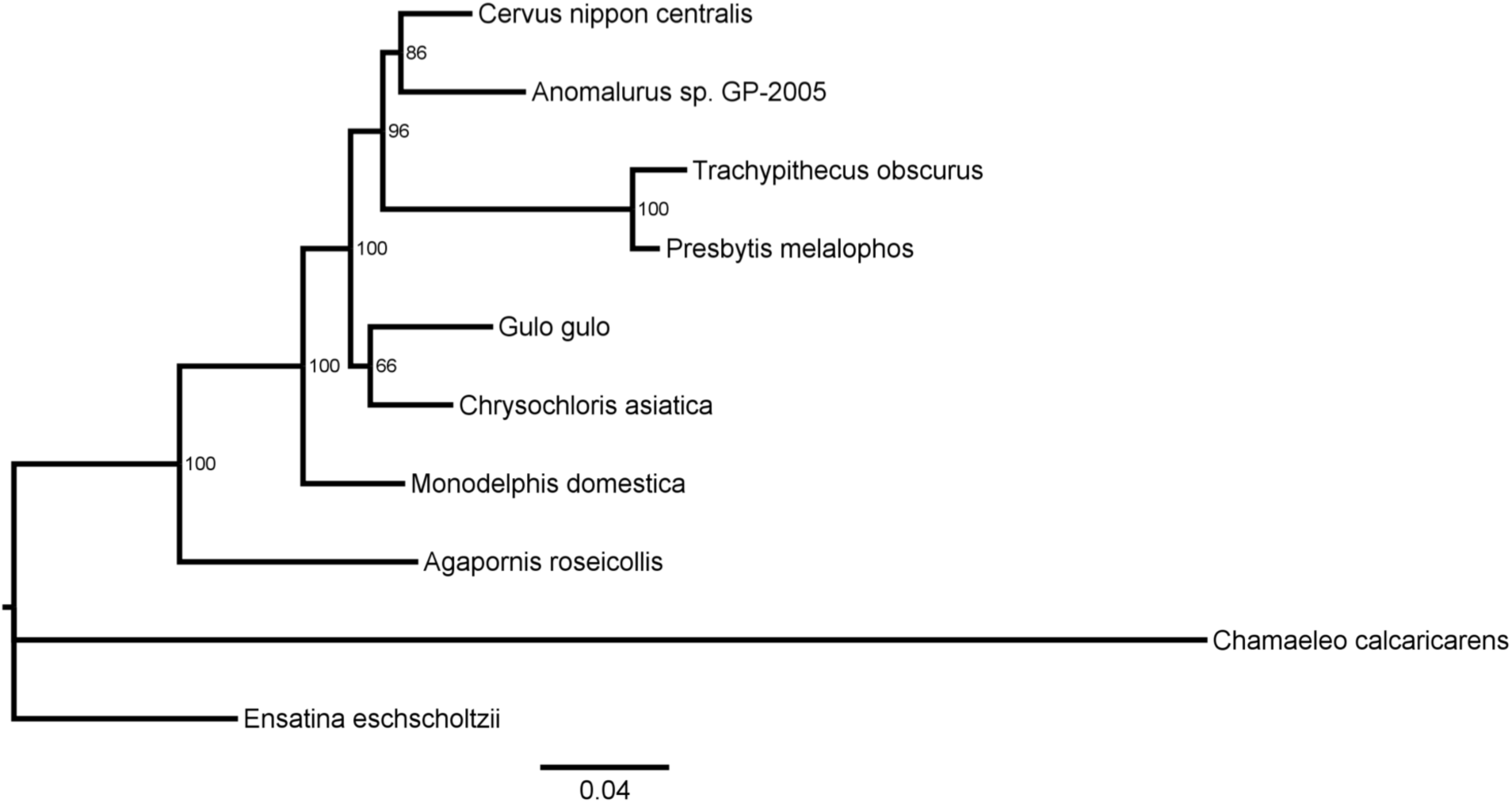
An example phylogenetic tree for the 10-taxon dataset. A tree for all protein coding regions in the mitochondrial genome is shown. Posterior probabilities are shown for the 495 amino acid Cytochrome C Oxidase subunit 1 (COI) alignment used in initial analyses. The posteriors were calculated using Mr. Bayes. Given the limited data used for the purpose of testing our method, this tree should not be interpreted as a true or species tree.

To understand the differences in topology sampling bias estimates obtained using different sample sizes, *f*, Markov katana was run with sample fractions ranging from 100% (495 sites) down to 20% (99 sites). The topology sampling biases for smaller *f* become somewhat more even, with the least frequent topologies about 10x more frequent for *f* = 20% than for *f*=100% (Fig. 3). At the same time, the number of topologies with sampling probabilities greater than 10^−6^ increased from 5,975 for *f*=100% to 21,198 for *f* = 20%. Predictably, comparisons of replicate sampling distribution runs indicated an increasing variance in estimated biases with decreasing probabilities in the runs (Fig. S2).

**Figure 3.**
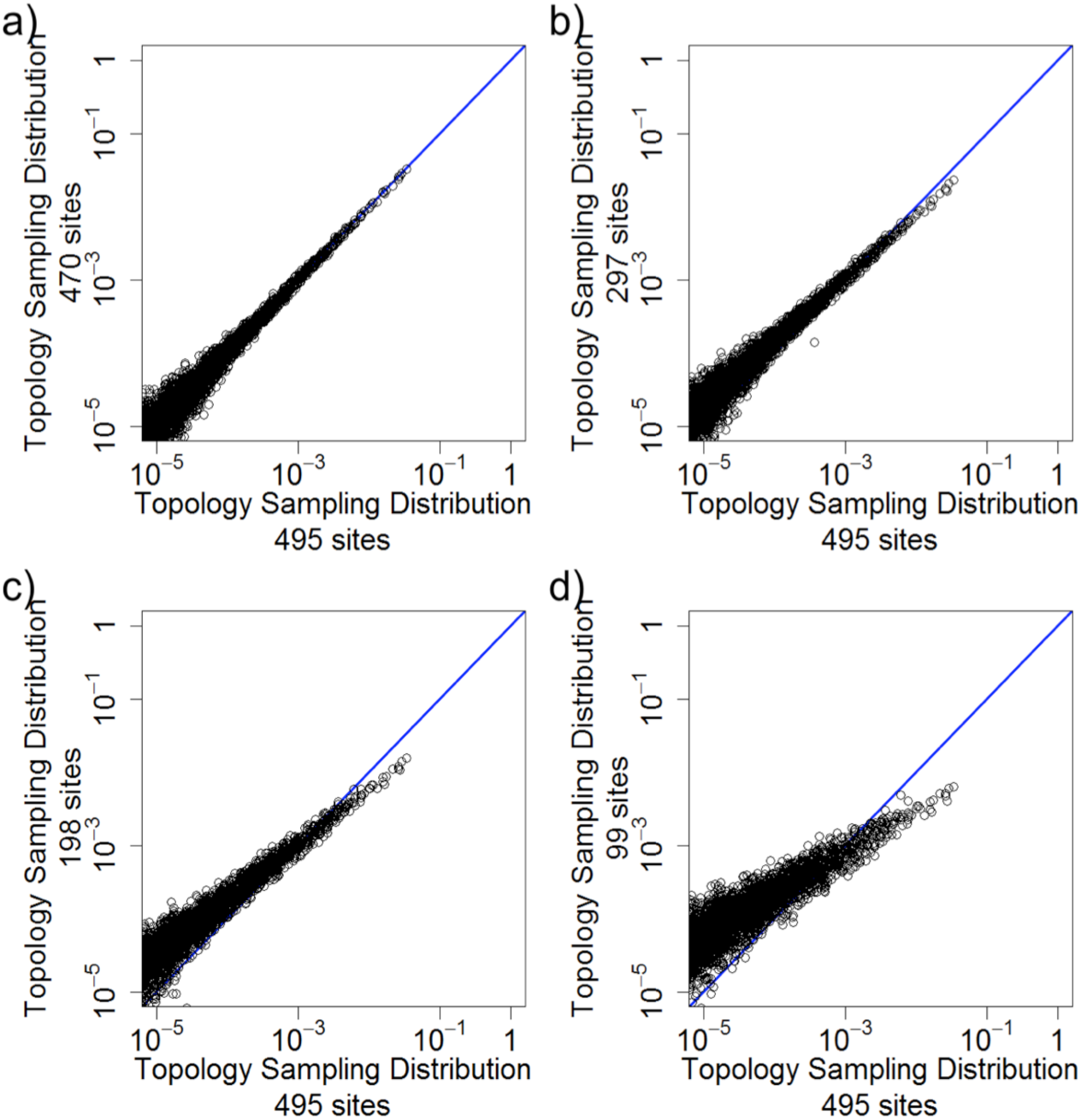
Relative sampling distributions estimations comparing different sample sizes. Sample size 1.0 sampling distribution versus the sampling distributions of sample size a) 0.95, b) 0.6, c) 0.4, d) 0.2. The blue line indicates where x and y values are equal. The sampling distributions were averaged over triplicate runs of 1,000,000 generations and used a jump size of 0.1.

The posterior correction (Equation 5) appears to work well across a broad range of sample sizes (Fig. 4). The corrected topology posteriors for sample fractions from 20% to 95% were all highly correlated with the topology posteriors for sample size 100%. It should be noted that the uncorrected posteriors are only slightly less correlated with each other than are the corrected posteriors (Fig.S3), meaning that the answer would have been similar without the correction. It is probably best to use the correction anyway, because in more complicated situations it may make more of a difference, and it is not too much trouble to obtain and is correct.

**Figure 4.**
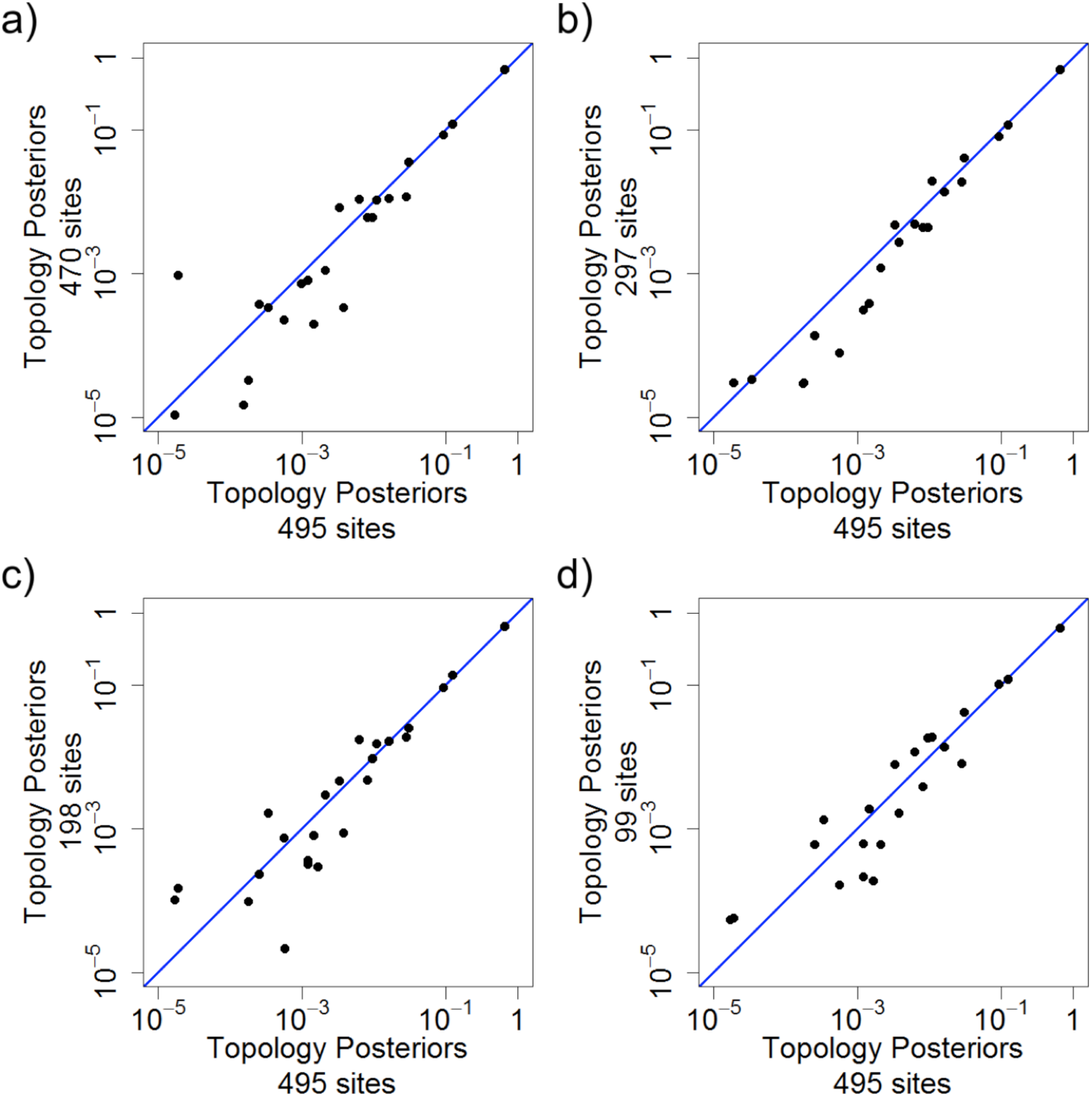
Corrected posteriors are linear over a wide range of sample sizes. Sample fraction 1.0 posteriordistribution versus the posterior distributions of sample fractions a) 0.95, b) 0.6, c) 0.4, and d) 0.2. The X = Y line is shown in blue. The uncorrected posteriors were averaged over triplicate runs of 50,000 generations and used a jump size of 0.1. The sampling distributions used for the correction were from Figure 3.

The corrected posterior estimates appear to be most noisy when the sampling distribution estimate is small and therefore poorly estimated. This is not entirely surprising given that the sampling distribution is in the denominator. Because the sampling distribution calculations are computationally inexpensive (they do not require a likelihood calculation), it is possible to obtain a couple orders of magnitude more data for them than for the uncorrected posterior estimates. While estimating the sampling distribution more precisely is important for the correction, many of the trees examined have topologies that are not found in the posterior. A potential means to increase accuracy of relevant topologies in the sampling distribution is to limit the sampling prior chains to those topologies seen in the uncorrected posterior.

### Effect of Sample Fraction and Jump Size on the Markov Chain

Although the posterior estimates were comparable for all sample sizes, it is still worthwhile to consider the effect of both sample size and jump size on the mixing efficiency of the Markov chain. For a range of conditions considered, the acceptance probability for Markov chain proposal varied from 10% to 90% (Fig. 5). We chose 100% sample size bootstraps for the NJ proposals along with a jump size of 50 (10%) as standard reference conditions, which had acceptance probabilities of about 30%.

**Figure 5.**
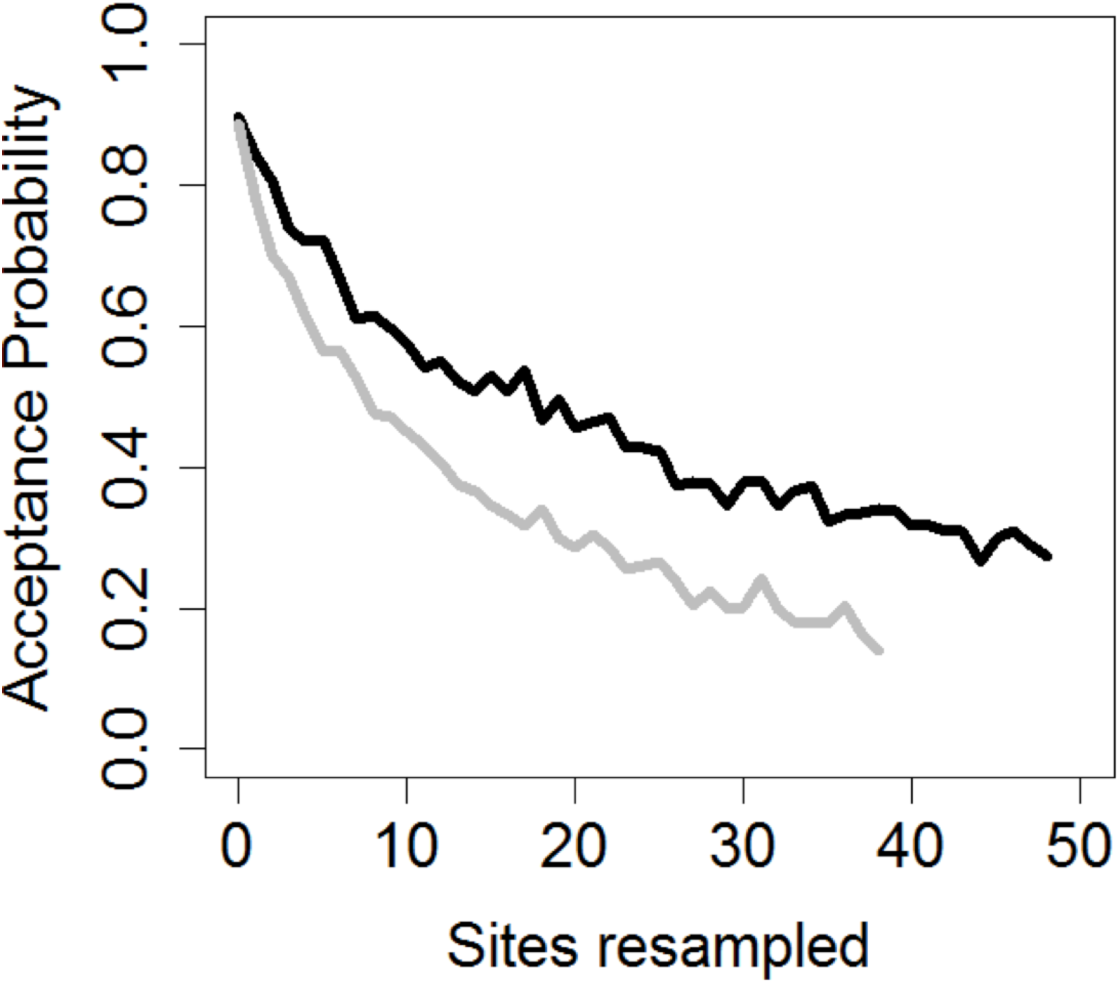
Acceptance probability for number of sites resampled (jump size). The average acceptance probabilities are shown for proposals that resampled different numbers of sites. For the dark line, the bootstrapped sample fraction for neighbor-joining (NJ) proposals was 100% and for the grey line it was 80%, both from the 495-amino acid 10-taxon dataset. Acceptance probabilities were determined by the average of 3 independent 50,000 generation runs of Markov katana.

We also considered the effect of jump size on both the sampling distribution and the uncorrected posterior Markov chain estimates. For the initial sampling distribution estimation procedure, the most well-mixed chain is of course the one with independent bootstraps (*j*=100%), but the chain also mixes well with lower jump sizes. It is necessary to have smaller jump sizes because a high proportion (99.9%) of the random samples are not in the uncorrected posterior topology set. For this analysis, the optimal jump size was j=0.85. This result did not differ much for a range of sample sizes. Although differing in detail, the jump size analysis for the uncorrected posterior had similar results to the biased sampling prior analysis.

Acceptance probabilities varied widely depending on jump size. In general, 5-10 sites appears to be a minimum, and 50 sites is probably a maximum. With smaller sample sizes (e.g., 80% shown here), the jump size is a larger proportion of the sample and reduces the acceptance probability more rapidly. Jump sizes bigger than 50 have somewhat greater probability of making large jumps in topology space (Fig. 6), at the cost of reduced probabilities of jumps tothe same topology due to lower acceptance probabilities.

**Figure 6.**
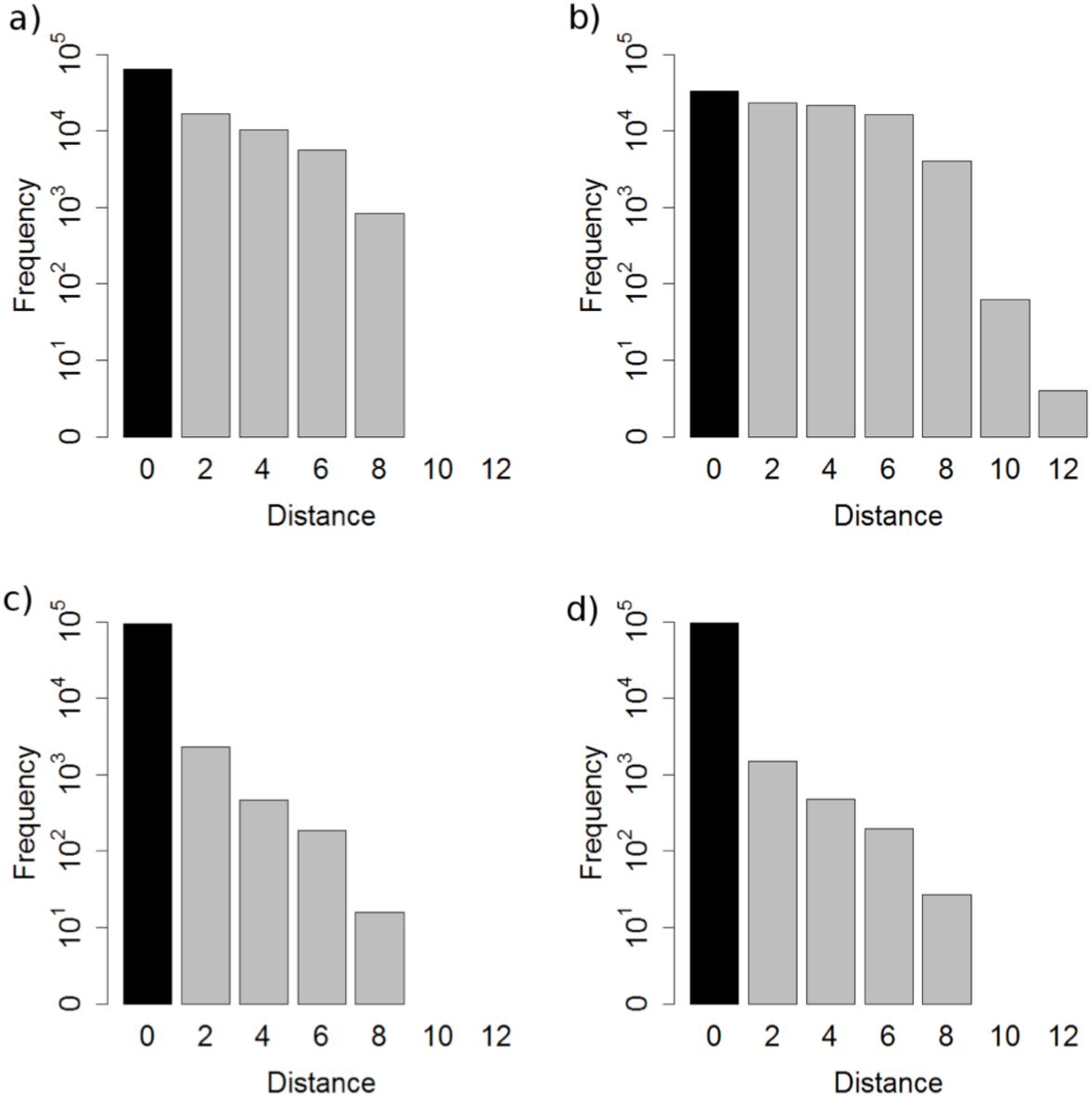
The effect of varying jump size. Increasing the jump size in general increases the average Robinson–Foulds (RF) distance between jumps and proposals. Parts a) and b) show the distance between trees in jumps for jump size 0.1 and 0.7 respectively. Parts c) and d) show the distance between trees in proposals for jump size 0.1 and 0.7 respectively. Rejected jumps are considered jumps of distance 0.

### The Structure of Tree Space

Figure 7 shows a network representation of the 12 tree topologies that had a posterior of 0.001 or higher (see Table 2). The size of the node represents the relative posterior of the topology, and the edges of the graph indicate NNI distances of one or two between the tree topologies. The tree topology space of this test data was clearly divided into two clusters of trees shown by the intragroup connections and the few intergroup connections. Given the connectivity of the network, other tree topology sampling procedures may have difficulty jumping between groups.

**Figure 7.**
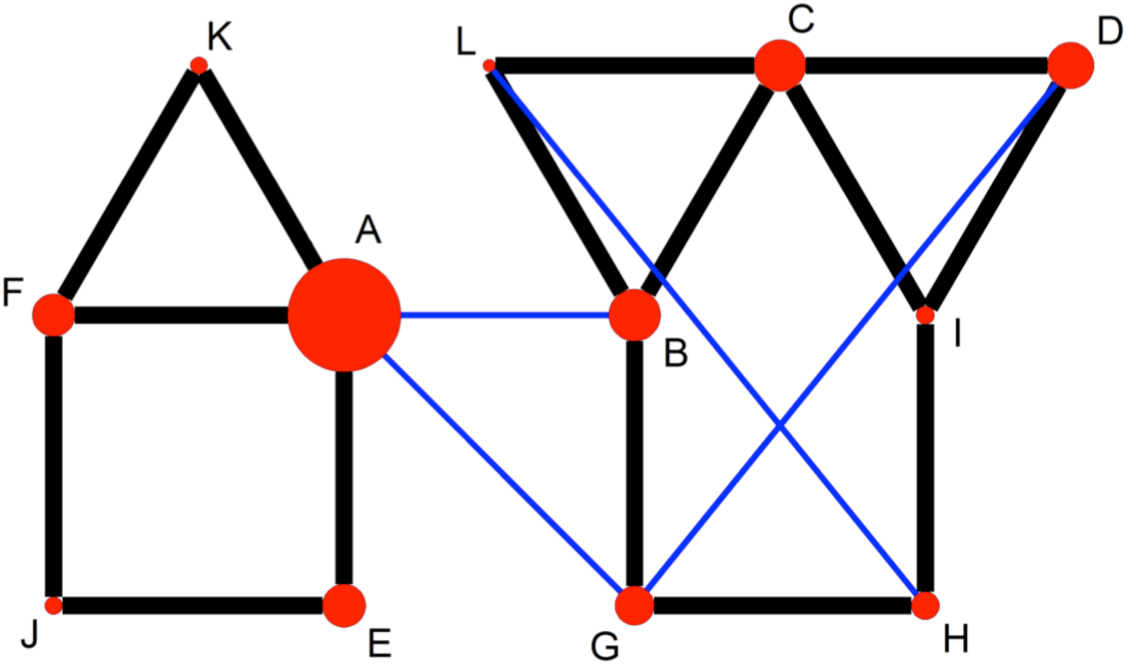
Tree distance network. This figure shows the distances between different tree topologies (A-L). The size of the circle shows the relative posterior probability. Black and blue lines indicate distances of 1 and 2 nearest neighbor interchange (NNI) (RF distance of 2 and 4), respectively. Topologies that are > 0.001 posterior probability.

**Table 2.**
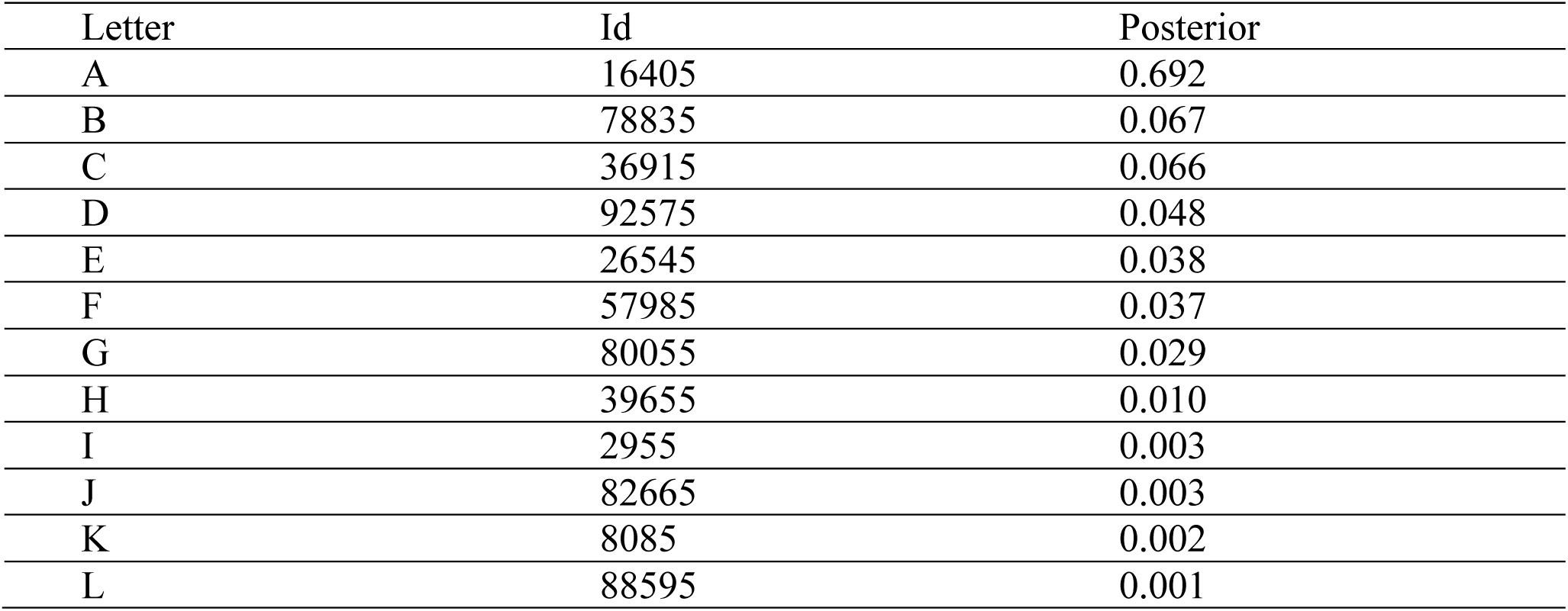
Posterior probability for topologies with substantial representation in the uncorrected posterior for the 10-taxon dataset. The topologies are labeled for reference in Figure 7.

| Letter | Id | Posterior |
| --- | --- | --- |
| A | 16405 | 0.692 |
| B | 78835 | 0.067 |
| C | 36915 | 0.066 |
| D | 92575 | 0.048 |
| E | 26545 | 0.038 |
| F | 57985 | 0.037 |
| G | 80055 | 0.029 |
| H | 39655 | 0.010 |
| I | 2955 | 0.003 |
| J | 82665 | 0.003 |
| K | 8085 | 0.002 |
| L | 88595 | 0.001 |

## DISCUSSION

We have demonstrated here that the Markov katana bootstrapping approach to phylogenetic tree searching can be a highly effective means for finding Bayesian posterior topologies and branches. It is able to take advantage of the speed of approximate distance-based methods to propose new trees, but retains the reliability of Bayesian methods. Many previous phylogenetic tree-search methods use the provided sequences for only the likelihood calculations, but Markov katana introduces a new way to explore tree space informed by thesequences. Including the sequence data in the tree search improves the fraction of high likelihood trees proposed and allows efficient jump proposals between even distant topologies.

For the 10-taxon dataset, the NJ algorithm is extremely fast, and the overall speed of the MK computation was limited by the likelihood calculations. As the number of taxa grows beyond ~200, the NJ algorithm slows dramatically and dominates computation times (data not shown). This could be alleviated using fast heuristic NJ algorithms or external programs such as RapidNJ that are optimized for large alignments (Simonsen 2008). Our current implementation calculates the distance contribution of each site only once and so is not hindered by the complexity of the distance measure. We did not see a great difference in the proposal bias for the two distance measures we compared, but further exploration of the performance of alternative distances may in some cases be warranted.

We used PAML for the likelihood calculations, but any program that computes likelihoods could potentially be used. The simplicity and adjustability of the approach means that it could be easily incorporated into existing sequence analysis packages (e.g., MrBayes, PAUP*, HyPhy, and PAML (Ronquist 2003; Swofford 2003; Pond 2005; Yang 2007)). We used a Perl script to implement the MK algorithm and demonstrate the method as simply as possible, but we expect that MK can be easily integrated directly into existing programs, which would then undoubtedly be much faster. We did not see the benefit in constructing a new likelihood program from scratch, although we believe the methodology would interact well with our existing context-dependent Bayesian analysis program, *PLEX* (de Koning 2012).

## ACKNOWLEDGEMENTS

Thanks to Seena D. Shah, who contributed to early versions of coding on *MarkovKatana*. Weacknowledge the support of the National Institutes of Health (NIH; GM083127 and GM097251) to DDP.

---

**Figure S1.**
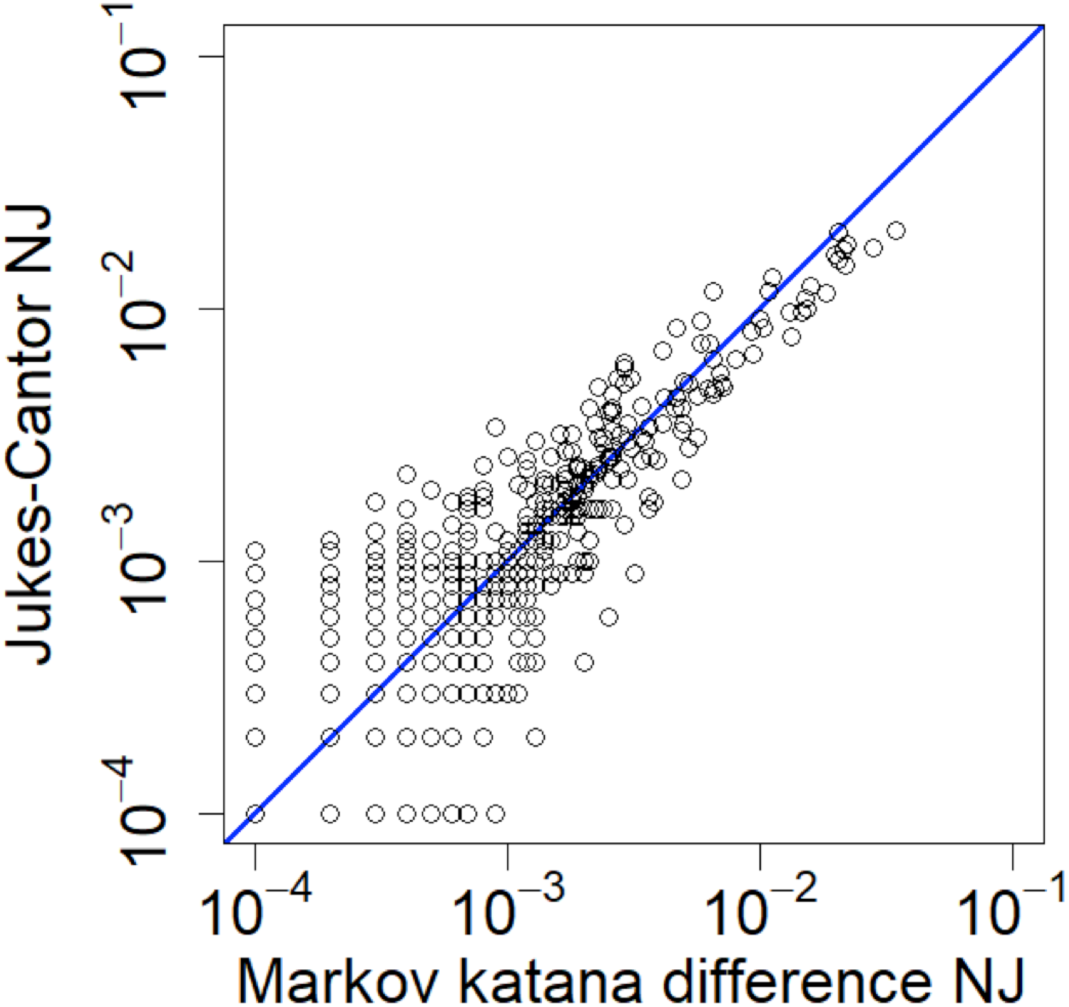
Supplementary. Comparing NJ implementations. Markov katana bootstrapped topology probabilities were compared with those from a different neighbor-joining program, RapidNJ. 10,000 bootstraps were run.

**Figure S2.**
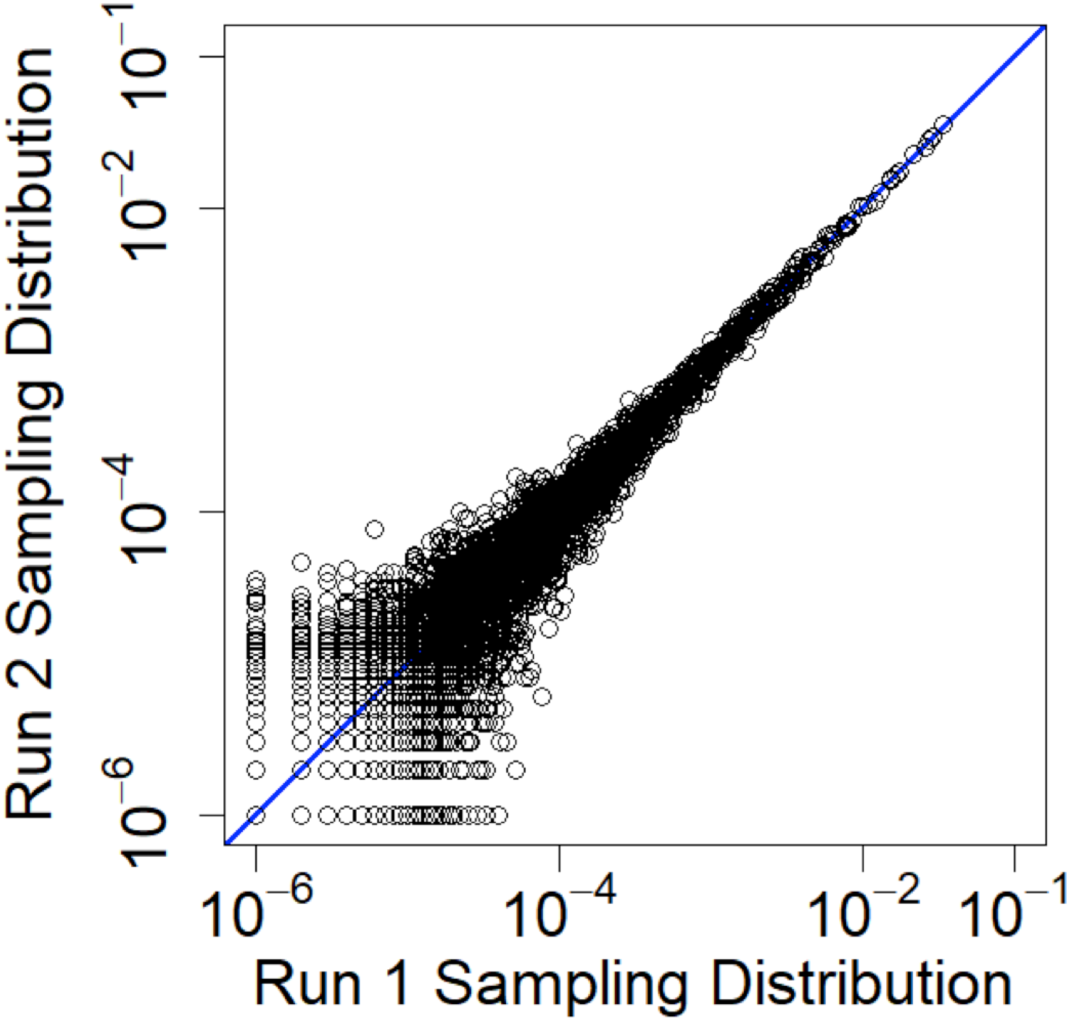
Supplementary. Variance of the sampling distribution among runs. The sampling distributions of two different runs with sample size 1.0 and jump size 0.1 and 1 million bootstraps are shown.

**Figure S3.**
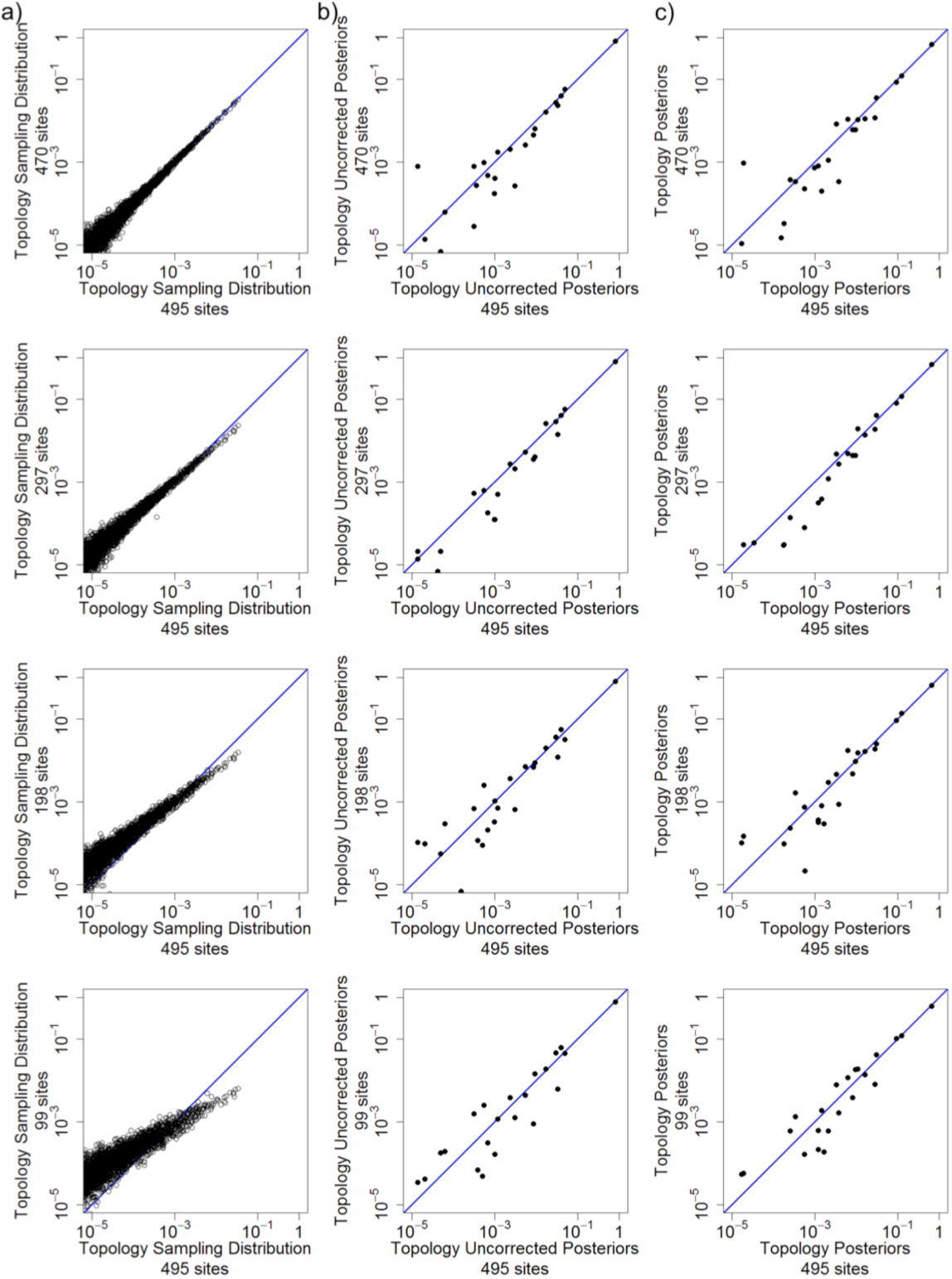
Supplementary. Comparing sampling distribution and posteriors across sample size. The sampling distribution (column a), uncorrected posteriors (column b) and corrected posteriors (column c) for 470 sites, 297 sites, 198 sites, and 99 sites, all versus 495 sites.

